# WAD: a wavelet-based linear programming method using L1-minimal reconstruction loss for accessible chromatin data deconvolution

**DOI:** 10.1101/2025.11.26.690776

**Authors:** Jeongho Chae, Benjamin McMichael, Terrence S. Furey

## Abstract

Bulk tissue-based accessible chromatin studies provide summary annotations across all cell types within the tissue. These annotations can be skewed by varying proportions of individual cell types, especially in the context of disease studies. Estimated sample specific cell-type proportions can be used to mitigate effects of this variability while also addressing whether there exist significant alterations in cell proportions under certain conditions, like disease. We present WAD (Wavelet-based Accessible chromatin Deconvolution), a principled framework for robust estimation of cell type composition of bulk accessible chromatin data such as from the ATAC-seq assay. To determine informative reference cell profiles from single-cell accessible chromatin studies, WAD leverages wavelet-based denoising to suppress stochastic noise while preserving local chromatin continuity. Cell type proportion inference is reformulated as an *L*_1_-minimal linear programming problem, enabling scalable and interpretable solutions. Across 700 *in silico* pseudo-bulk mixtures generated from single-cell data, WAD achieved a consistently lower mean absolute error (MAE) and higher concordance (*r >* 0.85) than existing machine learning-based methods. These results demonstrate that wavelet-based feature extraction provides a biologically grounded and computationally efficient approach to chromatin signal deconvolution. A complete implementation of WAD is available at https://github.com/chae-jh/WAD.

## 1 Introduction

Chromatin accessibility reflects the physical openness of regions of genomic DNA available for binding of regulatory proteins such as transcription factors (Klemm et al. 2019). It serves as a key layer of epigenetic regulation, along with other mechanisms such as DNA methylation and histone modifications, that contributes to gene expression levels and potentially disease mechanisms (Grandi et al. 2022, Buenrostro et al. 2013). The assay for transposase-accessible chromatin using sequencing (ATAC-seq) has become the standard approach for profiling genome-wide chromatin accessibility (Buenrostro et al. 2013, Klemm et al. 2019). Accessible chromatin studies in human tissues have increasingly shifted toward understanding molecular mechanisms at the cellular level. For bulk tissue studies, a critical challenge is distinguishing whether observed changes between samples with varying phenotypes, such as disease, arise from shifts in cell type composition or from changes within the same cell types (Avila Cobos et al. 2018). While single-cell sequencing technologies offer direct insights into the regulatory landscape at single-cell resolution, their low signal-to-noise ratio and substantial costs render them impractical for many biological studies (Tang et al. 2019, Lähnemann et al. 2020). To increase the utility and accuracy of bulk tissue analyses, computational deconvolution methods have emerged as a solution to the above challenges.

Estimated cell type-specific accessibility of deconvolved ATAC-seq data can be used to remove confounding effects of differing cell type proportions and identification of cell-specific regulatory changes in differential analyses, such as in disease, to enable a more accurate dissection of regulatory dynamics in complex tissues (Newman et al. 2019, Avila Cobos et al. 2018, Corces et al. 2017). Unlike gene expression, accessible chromatin data lacks a standard set of annotations requiring the data to also define these regions making feature selection for deconvolution difficult and computationally demanding (Ma et al. 2023). Growing numbers of single-cell ATAC-sequencing (scATAC-seq) datasets can be utilized to determine cell-specific annotations for deconvolution, but these datasets are generally sparse, noisy, and costly to generate at sufficient sequencing depth (Lähnemann et al. 2020, Chen et al. 2019). Further, current accessible chromatin deconvolution tools often rely on deep learning methods that require large-scale single-cell references for training and suffer from limited interpretability (Berson et al. 2023, Luo et al. 2025). Thus, there remains a need for new approaches that can robustly and interpretably estimate cell type proportions from bulk ATAC-seq data. To address these challenges, we introduce WAD (Wavelet-based Accessible chromatin Deconvolution), a novel method for robust cell type deconvolution of bulk accessible chromatin data. WAD provides several features that contribute to efficiency and accuracy including:

- Wavelet-based denoising: Single-cell accessible chromatin profiles are sparse and noisy. WAD employs wavelet-domain shrinkage to suppress background fluctuations while preserving local continuity to determine accurate and informative cell type reference profiles.
- Linear programming formulation: WAD reformulates the deconvolution problem into a linear program, enabling efficient scaling to tens of millions of peaks while leveraging solver-optimized sparse matrix operations.
- Robust to limited reference data: WAD remains effective even with low-coverage single-cell reference data thereby reducing reliance on large-scale datasets.
- Assumption-free applicability: WAD operates without tissue-specific priors or heuristic biological assumptions, enabling direct application across heterogeneous tissues, species, and sequencing platforms.

We show that WAD performs well compared to two existing machine learning deconvolution methods, Cellformer and DECA, and with greater user ease of use.

## 2 Methods

### 2.1 Overview of the WAD Framework

The WAD (Wavelet-based Accessible chromatin Deconvolution) pipeline, as illustrated in Figure 1, consists of two major steps: (i) construction of cell type chromatin accessibility profiles using single-cell data that includes wavelet-based denoising to suppress high-frequency background noise while preserving biologically informative chromatin structure; and (ii) deconvolution of bulk accessible chromatin data using a sparse linear programming formulation that minimizes differences between the bulk and deconvolved data using a L1-norm objective. In contrast to deep learning-based approaches, WAD is scalable to tens of millions of peaks, requires only a modest number of single-cell samples to construct a reliable cell-type profile, and yields interpretable coefficients directly corresponding to cell-type proportions.

**Figure 1.**
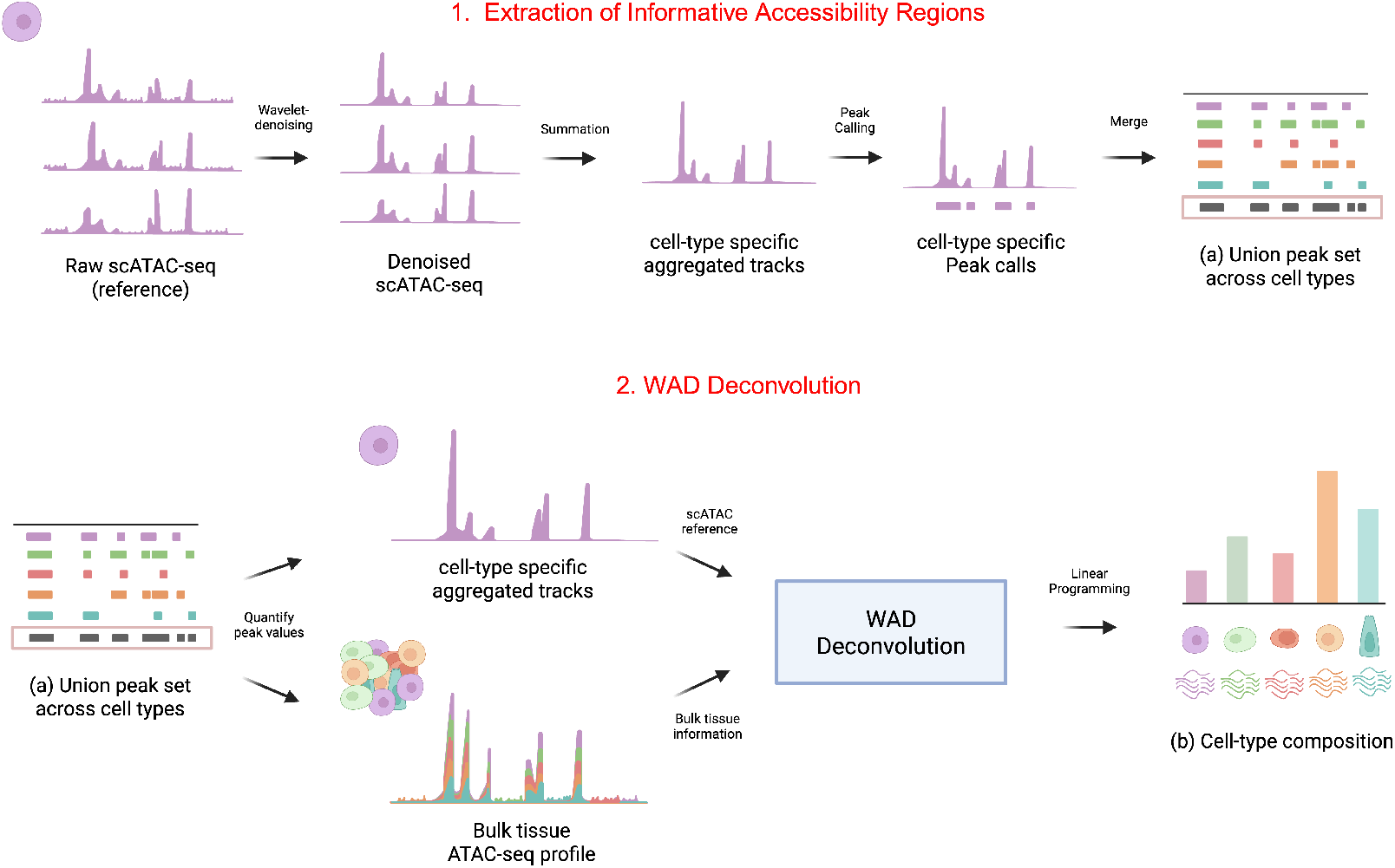
Overview of the WAD pipeline. (1) Extraction of informative accessibility regions: raw scATAC-seq profiles are denoised by wavelet transform, aggregated by cell type, and subjected to peak calling to construct a union peak set across cell types (a). (2) WAD deconvolution: bulk ATAC-seq profiles are quantified on this union peak set, and cell type composition is estimated (b) via linear programming.

### 2.2 WAD Stage 1: Construction of reference peak annotations

In the first step, WAD takes as input single-cell chromatin accessibility data, ideally from the same tissue being deconvolved. There are two main steps in the construction of the cell-specific profiles:

#### Wavelet modeling of single cell data

Wavelet transformation provides a compact time-frequency representation that is particularly well-suited for suppressing noise in sparse genomic signals, offering greater adaptivity than Fourier or other classical transformations. Single-cell chromatin accessibility data are often sparse and noisy with tens of millions of candidate peaks leading to an overdetermined system. Therefore, effective denoising is essential, but the continuity of the signal must be preserved for accurate peak detection (Hamilton and Furey 2023).

We explored the Daubechies (db) family as the wavelet basis function since it achieves the shortest support length given the number of vanishing moments, ensuring high local accuracy (Mallat 2009, Luo and Zhang 2012). We found that db4, with four vanishing moments, provided a balance between compact support and smoothness (Luo and Zhang 2012). Given functions *ϕ* and *ψ* as the scaling and wavelet basis functions, respectively, and the index *m* as the translation (position) of coefficients within each scale (Mallat 2009), reference signals were decomposed into *L* levels using db4 according to:

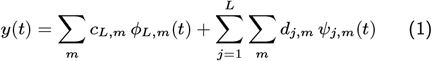

where *c*_*L,m*_ are approximation coefficients capturing the low-frequency structure, and *d*_*j,m*_ are detailed coefficients representing high-frequency variations of the signal.

Noise variance was estimated from the finest-scale detail coefficients using the median absolute deviation (MAD) estimator (Hampel 1974, Donoho and Johnstone 1994, Mallat 2009, Luo and Zhang 2012):

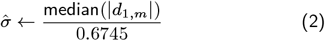

and the universal threshold was then defined as Donoho and Johnstone (1994) :

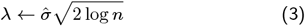

where *n* denotes the number of sampled points (peaks). Compared with SURE or minimax thresholds, the universal threshold is stricter, which is advantageous under an overdetermined state and noisy peak landscapes, ensuring stronger noise suppression (Mallat 2009, Luo and Zhang 2012). Finally, to avoid artificial discontinuities, we applied soft-thresholding rather than hard-thresholding, as introduced previously Donoho and Johnstone (1994):

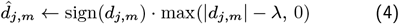

This overall approach suppresses noise while preserving local continuities necessary for peak calling. The complete denoising procedure is summarized in Algorithm 1.

##### Algorithm 1

Wavelet denoising with soft-thresholding.

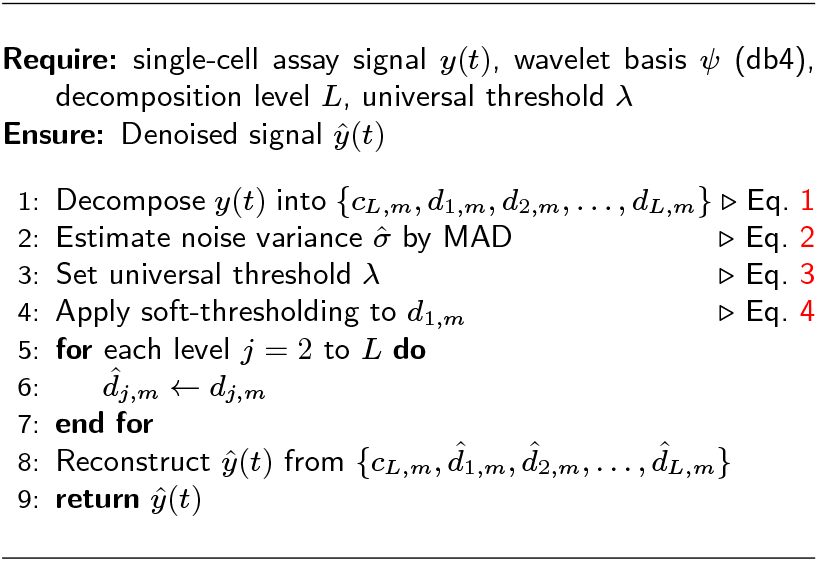

#### Reference peak set construction

Wavelet-denoised single-cell profiles were summed within each cell type to generate cell type-specific signal annotations. Cell type annotations were segmented into fixed genomic bins of 50 base pairs (bp). Genomic bins where more than 75% of the samples showed zero signal in the interval were removed ensuring only regions with informative coverage were considered.

Cell type-specific peak sets were identified using ROCCO (Hamilton and Furey 2023) with default settings except the budget parameter *b* that was fixed at 3% reflecting our expectation of the accessible proportion of the genome. A non-redundant final reference peak set was generated as the simple merged union of all cell type peaks.

### 2.3 WAD Stage 2: Deconvolution model and application

In the second stage, the reference peak set is used to define profiles for each cell type. These are used to estimate cell proportions in bulk tissue accessible chromatin data.

#### Reference profile matrix and bulk tissue input vector

Let *n* denote the number of reference peaks and *k* the number of cell types. The cell type-specific peak matrix *X* ∈ ℝ^*n*×*k*^ represents the collection of *k* cell type profiles, where each column is a cell type, each row is a reference peak genomic interval, and each *x*_*i,j*_ is the wavelet-denoised, aggregated signal in peak *i* for cell type *j*. The input bulk tissue data to deconvolve is represented as a vector *Y* ∈ ℝ^*n*×1^, where each element *y*_*i*_ denotes the raw read count observed in the genomic interval for peak *i*.

#### Deconvolution algorithm

The calculated cell type proportion vector is defined as *β* ∈ ℝ^*k*×1^, where each *β*_***j***_ represents the relative contribution of the cell type *j* to the bulk tissue accessibility signal. The deconvolution problem is formulated as:

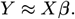

where the linear combination of cell-type-specific accessibility profiles in *X*, weighted by proportions in *β*, approximates the observed bulk profile *Y*. An overview of the matrix representation is shown in Figure 2, where the bulk profile is decomposed into cell-type-specific contributions via linear programming. While this matrix formulation provides a natural representation of the deconvolution problem, solving it directly is infeasible due to the overdetermined nature of the system (*n* ≫ *k*) and presence of substantial noise in accessible chromatin data.

**Figure 2.**
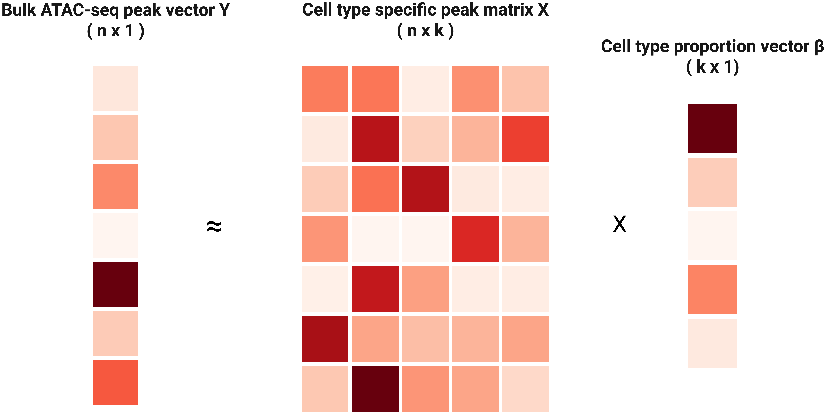
Matrix formulation of WAD deconvolution. The bulk ATAC-seq profile *Y* (left) is modeled as a linear combination of the cell-type-specific peak matrix *X* (middle) and the cell type proportion vector *β* (right). Each column of *X* represents a cell type, each row corresponds to a genomic interval(peak), and *β* captures the relative contributions of each cell type.

To overcome these challenges, we reformulate the deconvolution problem as an optimization task. The residual vector between observed bulk accessibility profile and reference-based reconstructed profile is defined as:

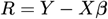

where *R* ∈ ℝ^*n*×1^ and each entry *R*_*i*_ represents the residual in the genomic interval for peak *i* between the bulk signal *y*_*i*_ and the predicted signal, and 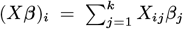 is the weighted sum of the cell-type reference peak signals. Robust estimation of *β* corresponds to minimizing the magnitude of these residuals. Given the susceptibility of accessible chromatin signals to noise, we adopted an L1-norm objective, denoted *f*_*COP*_ (*β*), which is well known to provide robustness against outliers and to promote sparse error distributions:

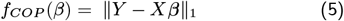

This objective is convex, ensuring that any local optimum is also a global optimum (Boyd and Vandenberghe 2004). To further enable efficient computation, we introduce a non-negative slack variable *s*_*i*_ that bounds the absolute residuals, a classical reformulation that converts *ℓ*_1_ regression into a linear program (Bertsimas and Tsitsiklis 1997):

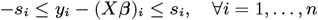

Substituting residuals with *s*_*i*_ transforms the deconvolution problem into a linear program (LP):

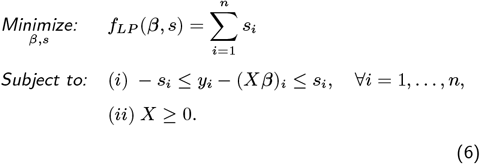

Linear programming, a subclass of convex optimization, guarantees polynomial-time solvability and is particularly efficient for large-scale sparse problems (Boyd and Vandenberghe 2004). Leveraging these properties, linear programming is efficiently solved using MOSEK (MOSEK ApS. 2023), a solver optimized for large-scale sparse LPs, that has been validated across numerous optimization-focused studies.

### 2.4 Liver tissue single-cell ATAC-seq data

Aligned single-cell ATAC-seq data from 39 human liver donors previously analyzed in our lab were used for evaluations (Alkhawaja et al. 2025). Cells were grouped into six major hepatic cell types (Cholangiocytes, Hepatocytes, Kupffer cells, Liver Sinusoidal Endothelial Cells (LSEC), Mesenchymal cells and NK-T cells) (see Supplementary Materials §3). To ensure high signal-to-noise ratio in the reference peak construction, we selected the ten samples exhibiting the highest transcription start site (TSS) enrichment scores for reference matrix generation. The remaining 29 samples were used for pseudo-bulk mixture construction and benchmarking analyses.

### 2.5 *In silico* pseudo bulk ATAC-seq construction

To evaluate cell type deconvolution under controlled yet realistic conditions, we constructed synthetic *in silico* pseudo-bulk ATAC-seq samples varying their composition along four experimental design characteristics:

a. Evenness of cell proportions: Cell type proportions were drawn from a Dirichlet prior with concentration parameter *α* ∈ {1, 2, 5}. Smaller *α* values produce sparser, more imbalanced mixtures of cells, whereas larger *α* yield more uniform compositions. Default *α* = 2.
b. Inclusion of a rare cell type: A designated cell type was fixed at a small fraction (0.5, 1, 2, 5%) of the mixture. The remaining cell type proportions were sampled from a Dirichlet distribution and renormalized so that the sum of all cell proportions was 100%. Default = no rare cell type.
c. Effective input sample number, *N*_eff_: The number of input samples from which aligned sequences were sampled was modulated by choosing either a small (4-8), medium (12-16), or large (20-24) input samples. The fraction of reads taken from each input sample was based on a Dirichlet distribution, with no input sample contributing more than 50% of the total reads to the pseudo-bulk sample. Increasing the number of input samples considered affects the degree of pseudo-bulk sample heterogeneity. Default *k* = medium number of input samples (12-16).
d. Sequencing depth, *K*_tot_: Given the above constraints, a target of 10, 20, 50, or 100 million reads (corresponding to 5, 10, 25, and 50 million read pairs, respectively) was set for each pseudo-bulk sample. Default depth = 50M reads.

#### Generation of ***in silico*** data

Cell type proportions **w** = (*w*_1_, …, *w*_6_) for a given pseudo-bulk sample were first determined using a 5-simplex Dirichlet prior (Murphy 2012). By default, we used a symmetric parameterization:

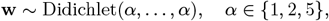

To confirm the evenness of the cell proportions generated, we calculated the Shannon entropy

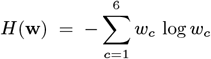

with the natural logarithm (units: nats) (Shannon 1948). Larger *H* indicates a more even mixture and attains a maximum at the uniform composition 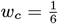, for which

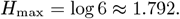

To generate *K*_tot_ paired-end reads for a pseudo-bulk sample, we calculated *K*_*c*_ = *K*_tot_ *w*_*c*_ for each cell type *c* and converted this to the expected number of contributing cells using empirically estimated median reads-per-cell *μ*_*c*_ from the liver scATAC-seq barcode catalog:

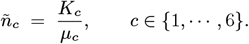

Cell counts **n** = (*n*_1_,. .., *n*_6_) were rounded to an integer using largest remainder method (LRM) to maintain the total sample read count to within a rounding error while minimizing distortion of the target mixture. For each cell type, we then designated a certain number of *k* input samples from predefined number ranges from which actual aligned reads will be used in the pseudo-bulk sample:

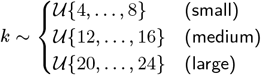

To measure the effective number of resulting input samples, we calculated *N*_eff_: (Laakso and Taagepera 1979):

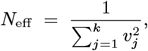

where **v** = (*v*_1_, …, *v*_*k*_) reflects the proportion of reads from each input sample drawn from a Dirichlet distribution Dirichlet(*α*_***1***_) (default *α*_sample_=2) with maximum *v*_*j*_ ≤ 0.5, meaning no more than 50% of the reads could come from a single input sample. Final cell counts used from each designated input sample were calculated as *n*_*c*.*j*_ = *n*_*c*_ *v*_*j*_ for each cell type *c* in input sample *j*, rounded to integers using LRM (Pukelsheim 2014).

Lastly, for each cell type *c* and input sample *j*, we sampled *n*_*c,j*_ distinct cell barcodes without replacement from the set of barcodes observed for each cell type in these data. Reads were then extracted from the liver scATAC-seq BAM files by barcode using <monospace>samtools</monospace>. We excluded unmapped reads, secondary alignments, alignments that failed QC, duplicate reads, and supplementary alignments while requiring alignment mapping quality MAPQ ≥ 30. Cell type aligned reads to be used for a given pseudo-bulk sample were merged, coordinate-sorted and indexed. When necessary, reads were down-sampled according to:

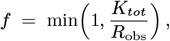

where *R*_obs_ denotes total merged reads and *R*_obs_ ≤ *K*_tot_ indicates no down-sampling.

### 2.6 Benchmarking with existing methods

To assess performance, we benchmarked against two other ATAC-seq deconvolution methods, Cellformer and DECA (Berson et al. 2023, Luo et al. 2025). Each tool was executed following the authors’ official GitHub implementations and default hyperparameter settings, substituting only the input datasets with our pseudo-bulk ATAC-seq mixtures.

## 3 Results

### 3.1 Generation of ***in silico*** pseudo-bulk datasets using liver scATAC-seq data

Evaluation of deconvolution methods requires datasets for which the cell composition is known. Using scATAC-seq data from human liver samples that consisted of six diverse cell populations, we generated 700 *in silico* pseudo-bulk ATAC-seq samples. These pseudo-bulk samples were constructed by varying four key characteristics: evenness of the proportion of cells from each cell type, presence of a rare cell type (<5%), the number of reference samples that contributed to the pseudo-bulk sample data, and sequencing depth (see Methods §3.5 for full procedure). We verified that the generated samples accurately adhered to these desired characteristics (see Supplementary Materials §1).

To ensure that these design factors did not confound sample quality of the *in silico* dataset, we evaluated the resulting fraction of reads in peaks (FRiP) and transcription start site enrichment score (TSS), two standard accessible chromatin quality metrics, across our variable sample characteristics. Median FRiP values were stable around 0.17 (5-95% range: 0.152 - 0.192), while median TSS enrichment scores were consistently results confirm that the ∼3.7 across all conditions. These pseudo-bulk samples and cohorts preserved uniform library complexity and promoter-centered signal quality, independent of the experimental factors being tested.

### 3.2 Performance evaluation of WAD across wavelet decomposition levels

We first used the liver pseudo-bulk data to evaluate WAD deconvolution performance. In our evaluations, we realized that a key factor in the first step, the generation of reference cell type profiles, is the choice of the wavelet decomposition level, *L*. Intuitively, this controls the granularity of signal separation during denoising (see Algorithm 1 and Methods §3.2). Low wavelet levels yield a coarser representation of the accessibility signal suppressing high-frequency signal noise but potentially missing finer details of the chromatin landscape. Conversely, high wavelet levels iteratively decompose the signal into multiple frequency bands, providing a smoother and more nuanced reconstruction of the accessibility signal that preserves subtle variations at the expense of weaker noise attenuation. Thus, varying *L* permits WAD to explore the trade-off between denoising strength and spatial precision in cell type chromatin accessibility profiles.

Therefore, we evaluated WAD using five different wavelet levels. Overall, WAD achieved strong concordance between predicted and true proportions for all levels with Pearson correlation coefficients (*r*) ranging from 0.87-0.96 across a wide range of predicted and observed proportion levels of hepatocytes (Figure 3). To more systematically evaluate performance, we used the mean absolute error (MAE) to compare across our four experimental design factors for each wavelet level.

**Figure 3.**
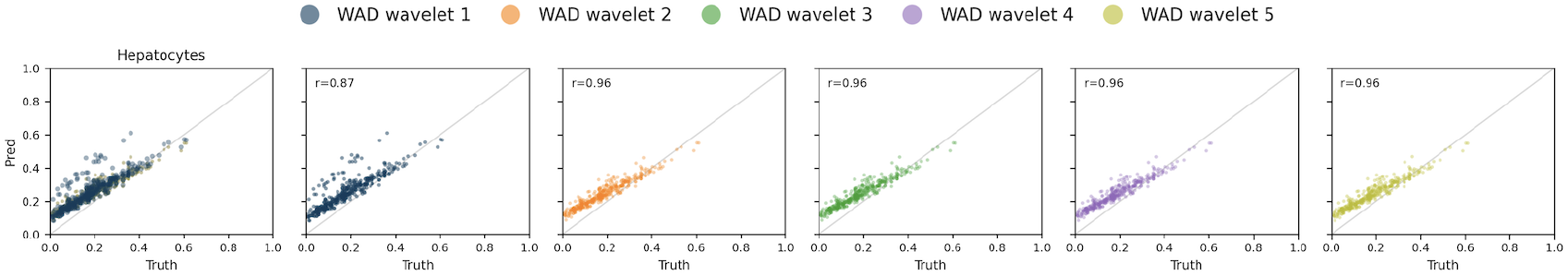
Truth-prediction scatter plots for Hepatocytes across WAD wavelet levels. Each point represents one *in silico* pseudo-bulk sample, with the *x*-axis showing ground-truth Hepatocyte proportion and the *y*-axis showing the predicted proportion. The leftmost panel overlays all wavelet levels (1-5) for direct visual comparison, while the five panels on the right show individual wavelet levels separately. The grey diagonal line (45^°^) denotes perfect agreement between predicted and true proportions. The Pearson correlation coefficient (*r*) denotes concordance between predicted and true proportions across 700 pseudo-bulk realizations.

Quantitatively, level 1 achieved the lowest overall MAE (0.0615) and high concordance (*r* = 0.85 across all cell types), outperforming higher wavelet levels (MAE ≈ 0.0635-0.064) (see Supplementary Materials §2). Across experimental factors (Fig. 4a-d), wavelet 1 again consistently yielded the lowest MAE for mixture evenness (*α* = 1, 2, 5), rare-cell spike-ins (0.5-5%), and number of contributing samples (*N*_eff_ bins), indicating that stronger denoising at the first decomposition level effectively suppresses stochastic noise while retaining biologically informative structure. Interestingly, sequencing depth was the exception to this trend (Fig. 4b): under the lowest depth (10M read pairs), wavelet level 1 exhibited the highest MAE, whereas higher wavelet levels (2-5) maintained a smaller error. At low coverage, overly aggressive filtering in wavelet level 1 appears to attenuate legitimate peak signals, reducing contrast and impairing decomposition accuracy. However, beyond 20M reads, which is closer to the ENCODE recommended bulk ATAC-seq sequencing depth of approximately 50M reads, wavelet level 1 outperformed all other levels, suggesting that at sufficient depth, it is still the best choice. The overall behavior of wavelet level 1 reflects the adaptivity of this discrete wavelet transform, which captures coarse but stable accessibility domains that then improves regression conditioning during deconvolution. Such coarse-scale smoothing is particularly well-suited to ATAC-seq data, whose inherently sparse coverage and low signal-to-noise ratio benefit from reduced local variance in the accessibility signal.

**Figure 4.**
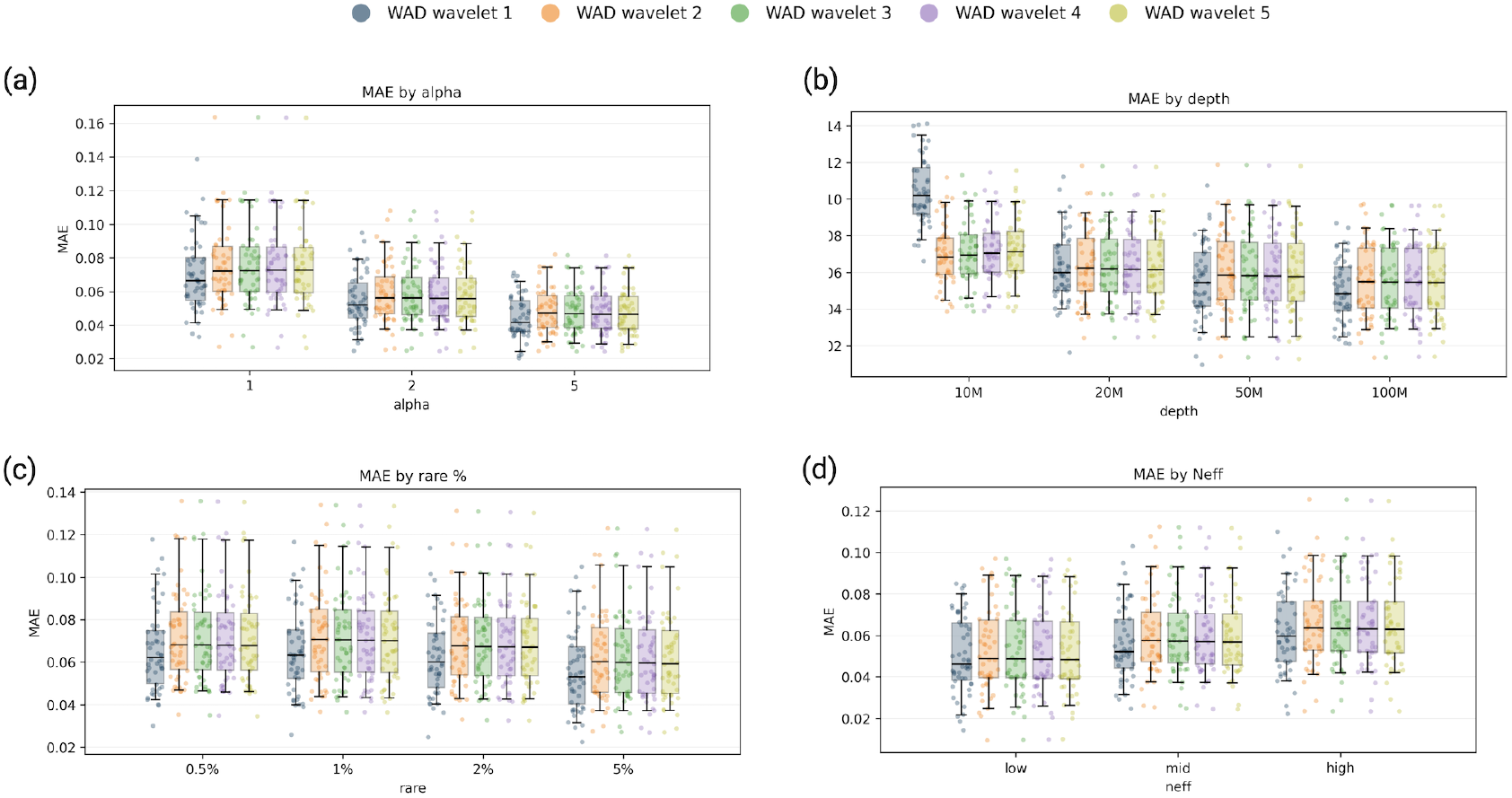
WAD deconvolution accuracy across design factors. Box-and-jitter plots compare five wavelet levels (colors in legend) for mean absolute error (MAE) computed per sample as the average 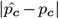 across six liver cell types. (a) MAE by Dirichlet concentration *α* ∈ {1, 2, 5}. (b) MAE by sequencing depth (10M, 20M, 50M, 100M unique reads). (c) MAE across rare NK-T spike-in levels (0.5%, 1%, 2%, 5%). (d) MAE by effective contributor count (*N*_eff_: low, mid, high).

We further investigated this depth-dependent effect in one cell type, hepatocytes (Figure 5), comparing pseudo-bulk samples restricted to 10M reads versus those with higher coverage. Although samples in both the low and high depth sets showed high correlation between the estimated and actual cell proportions (*r* = 0.96 − 0.97), the low-depth samples displayed greater dispersion from the diagonal, confirming that excessive denoising under sparse coverage may obscure weak but genuine accessibility signals. These results reinforce that the apparent lower performance of wavelet level 1 at 10M arises primarily from coverage-induced signal attenuation.

**Figure 5.**
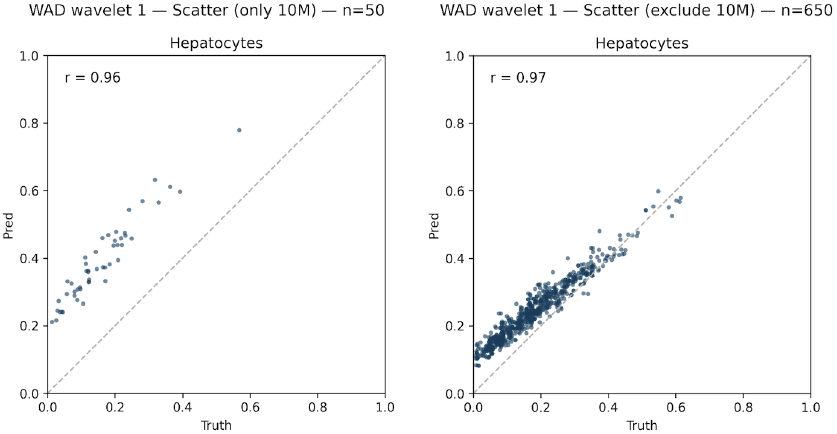
Effect of sequencing depth on WAD wavelet 1 prediction accuracy. Scatter plots show truth-versus-prediction concordance for Hepatocytes under the shallowest wavelet decomposition (level 1). The left panel includes only low-depth pseudo-bulks (10M read pairs, *n* = 50), while the right panel excludes them (*n* = 650).

Beyond accuracy, we evaluated the computational efficiency of WAD across wavelet decomposition levels (Table 1). All analyses were executed on CPU-only hardware using the *mosek* solver for the linear programming (LP) stage. Despite its mathematical formulation, WAD showed high computational efficiency: the LP solver, which is responsible for estimating cell-type proportions, dominated the runtime, averaging roughly 2 minutes per bulk sample. Wavelet denoising and ROCCO peak calling, although more computationally intensive in isolation, are performed only when building the reference profiles and thus are not affected by increased numbers of bulk samples to deconvolve. Consequently, the full WAD pipeline completes end-to-end deconvolution using only CPUs, without the need for GPUs or model retraining. This stands in sharp contrast to many deep learning-based frameworks, whose runtimes depend heavily on specialized GPU hardware and extensive training overhead.

**Table 1.** WAD deconvolution runtime (LP) across wavelet levels. Each value represents the mean per-sample wall time (in seconds) for the linear-programming deconvolution step, computed over 700 pseudo-bulk samples per wavelet level (CPU-only).

| Wavelet level | Runtime (s) |
| --- | --- |
| 1 | 161.65 |
| 2 | 90.14 |
| 3 | 100.54 |
| 4 | 115.06 |
| 5 | 44.88 |

Collectively, these results demonstrate that: (i) WAD maintains high overall fidelity (*r >* 0.9) across most choices of wavelet level; (ii) wavelet level 1 provides the best balance between denoising and signal retention under realistic sequencing conditions (reads ≥ 20*M*); and (iii) higher wavelet levels introduce marginal smoothing that slightly degrades sensitivity to rare cell types and low-depth signals. Accordingly, wavelet level 1 was adopted as the default for all subsequent analyses.

### 3.3 Comparison with machine learning-based deconvolution methods

We compared WAD against two state-of-the-art machine learning-based deconvolution tools, *Cellformer* and *DECA*, using the 700 generated liver pseudo-bulk samples (Berson et al. 2023, Luo et al. 2025). We found that WAD consistently exhibited the lowest error rates and highest concordance between predicted and true cell-type proportions (Figure 6). Across all samples, the mean/median MAE was 0.062/0.058 for WAD versus 0.077/0.074 for DECA and 0.148/0.149 for Cellformer. We also calculated the mean/median root mean square error, RMSE, which showed a similar pattern (0.074/0.069 for WAD vs 0.097/0.093 for DECA and 0.177/0.178 for Cellformer). Concordance was also higher for WAD (mean Pearson *r* = 0.635, median 0.731) than for DECA (0.496, median 0.618), while Cellformer exhibited a near-zero negative mean (mean -0.028, median -0.053), indicating unstable cross-cell type agreement. Dispersion was also smallest for WAD, as the standard deviation of the MAE/RMSE across all pseudo-bulk samples was 0.023/0.029 for WAD compared to 0.029/0.040 for DECA and 0.035/0.039 for Cellformer. Together, these results show that WAD yielded lower error and higher correlation than both ML-based baselines over a broad range of mixture compositions.

**Figure 6.**
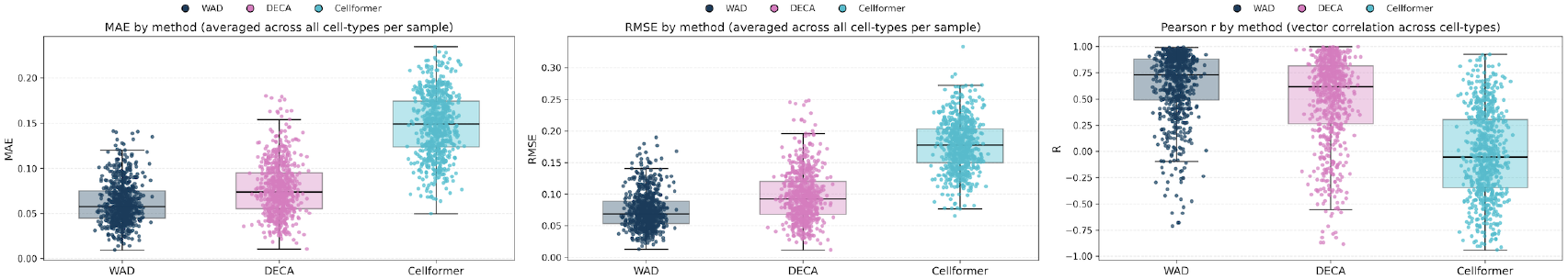
Cross-method comparison of deconvolution accuracy on *in silico* pseudo-bulk ATAC-seq mixtures. Box-and-jitter plots comparing three methods—WAD, DECA, and Cellformer—over mean MAE (left), RMSE (middle), and Pearson r (right).

When stratified by experimental design factors (Figure 7), WAD showed better overall performance across different levels of mixture evenness (*α*), when including rare cell types, and when varying the number of input samples used to create the pseudo-bulk sample (*N*_eff_). As shown in Fig. 7a, mean MAE steadily decreased with increasing mixture evenness for both WAD (*α* = 1 → 5, 0.0703 → 0.0444) and DECA (0.0877 → 0.0697) but did not vary substantially with Cellformer (∼ 0.15 across all levels). At the lowest sequencing depth (10M), the performance for all methods declined likely due to read sparsity, but the effect was most pronounced for WAD (MAE 0.104) owing to the strong denoising effect of wavelet 1 decomposition as shown above (Fig. 7b). However, at more typical sequencing depths of >20M reads, WAD showed the smallest error (MAE 0.063 at 20M, 0.055 at 50M, and 0.050 at 100M) compared to DECA and Cellformer. Across rare-cell spike-in fractions (Fig. 7c) and number of input samples (Fig 7d), WAD consistently showed the smallest mean and median MAE with reduced variability, whereas DECA showed intermediate accuracy and Cellformer performed worst across all metrics.

**Figure 7.**
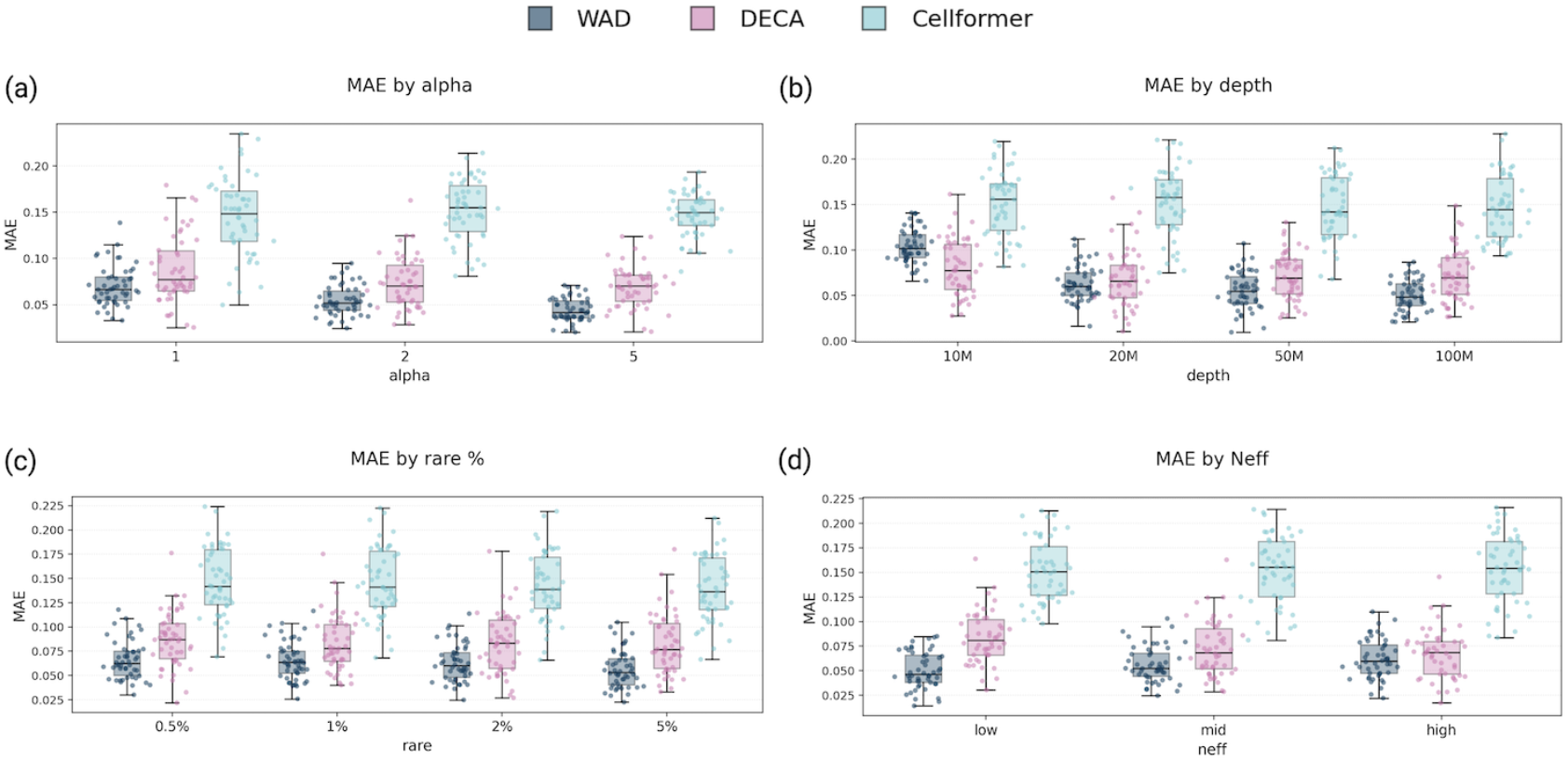
Head-to-head deconvolution accuracy across design factors. Box-and-jitter plots compare WAD (wavelet 1), DECA, and Cellformer on the same *in silico* pseudo-bulks. Each point is one pseudo-bulk realization. (a) MAE by Dirichlet concentration (*α* = 1, 2, 5). (b) MAE by sequencing depth (10/20/50/100M read pairs). (c) MAE by rare NK-T spike-in (0.5/1/2/5%). (d) MAE by donor diversity bin (*N*_eff_ : low/mid/high).

Together, these results indicate that WAD’s wavelet-based representation captures coarse but biologically meaningful chromatin patterns that enhance regression conditioning, especially under low signal-to-noise conditions characteristic of ATAC-seq data.

## 4 Discussion

The increasing availability of chromatin accessibility data across tissues and biological conditions highlights the need for interpretable and computationally efficient deconvolution methods. Here, we introduced WAD (Wavelet-based Accessible chromatin Deconvolution), a model-free method that leverages discrete wavelet transforms to separate biologically meaningful accessibility signals from stochastic background noise to accurately generate cell type reference models prior to deconvolution. Benchmarking analysis using 700 synthetic pseudo-bulk mixtures derived from 39 primary human liver donors, WAD consistently outperformed two state-of-the-art machine learning (ML) methods, DECA and Cellformer, achieving higher accuracy, stability, and reproducibility across diverse mixture conditions.

### Wavelet-based representation enhances signal fidelity in sparse chromatin data

ATAC-seq data are inherently sparse and noisy, especially at low sequencing depths and in cell types with limited chromatin accessibility. WAD directly addresses this challenge by decomposing single-cell signals into multi-resolution components, effectively isolating coarse-scale accessibility domains while suppressing high-frequency noise. The results across decomposition levels demonstrated that the first-level wavelet transform (level 1) provided the optimal balance between denoising and structural preservation: strong enough to stabilize regression, yet sensitive enough to retain biologically informative peaks. This denoising property may explain WAD’s decreased mean absolute error (MAE) and higher concordance compared to ML-based models, which may overfit stochastic fluctuations in sparse genomic regions.

### Performance under coverage constraints

While WAD achieved the lowest error under typical sequencing conditions (≥ 20*M* reads), performance was degraded at lower depths (10*M* reads) likely due to the aggressive noise suppression in low-level wavelet decompositions. This behavior reflects a general trade-off in deconvolution: at low signal-to-noise ratios, smoothing can attenuate true peaks, reducing contrast between cell-type-specific features. Nonetheless, this limitation is only seen at sequencing depths below what is typically obtained and far less than the ENCODE-recommended depth for bulk ATAC-seq (≈ 50M reads).

### Future directions

This wavelet-based paradigm underlying WAD could generalize to sequencing data generated for other regulatory annotations such as DNA methylation, histone modifications, or multi-omic ATAC-RNA-seq datasets. Importantly, because WAD does not rely on tissue-specific priors or biological assumptions, or retrained reference models, it can be directly applied to datasets from diverse organs, species, or experimental conditions with minimal adaptation. Integration with spatially resolved epigenomics or single-cell multiome data could further exploit the multiscale nature of wavelets to model cell state-specific chromatin landscapes. Additionally, combining WAD with probabilistic or Bayesian deconvolution models may lead to hybrid frameworks that retain interpretability while capturing nonlinear dependencies between diverse regulatory factors.

### Concluding remarks

Overall, WAD establishes a principled, interpretable, and computationally efficient method for chromatin accessibility deconvolution. By introducing wavelet denoising as a preprocessing step, WAD connects statistical signal processing and functional genomics, offering a scalable alternative to machine learning models. Its performance across diverse synthetic mixtures from real single-cell data supports the potential of signal-domain modeling to increase the precision of deconvolution in epigenomic research.

## Code Availability

The WAD framework (v1.0.0) was used to generate all results reported in this study. The source code is openly available at https://github.com/chae-jh/WAD. User documentation including API reference and usage tutorials is accessible at https://chae-jh.github.io/WAD.

