## Supplementary Materials for "WAD: a wavelet-based linear programming method using L1-minimal reconstruction loss for accessible chromatin data deconvolution"

#### 1 Validation of *in silico* pseudo-bulk data generation

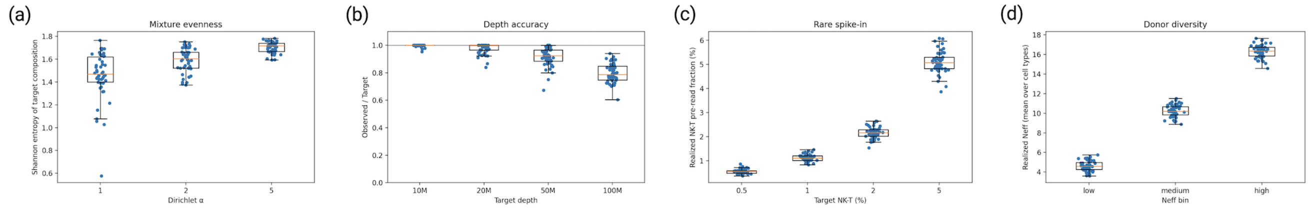


**Figure 1. Design validation of the *in silico* pseudo-bulk panel. (a)** Mixture evenness increases with the Dirichlet concentration parameter ($\alpha$); points are individual pseudo-bulks and boxes show interquartile range (IQR) with median (whiskers: $1.5\times$IQR; $n = 50$ per level). **(b)** Depth accuracy, reported as observed-to-target read ratio across target depths (10M, 20M, 50M, 100M); the grey line marks perfect tracking ($= 1.0$). **(c)** Rare NK–T spike-ins: realized pre-read fractions by target level (0.5%, 1%, 2%, 5%); small dispersion reflects integer cell selection, barcode availability, and filtering. **(d)** Donor diversity by bin (low/medium/high), summarized by realized $N_{\text{eff}} = 1/\sum_j v_j^2$ averaged across cell types. All panels use box-and-jitter plots; $n = 50$ pseudo-bulks per condition.

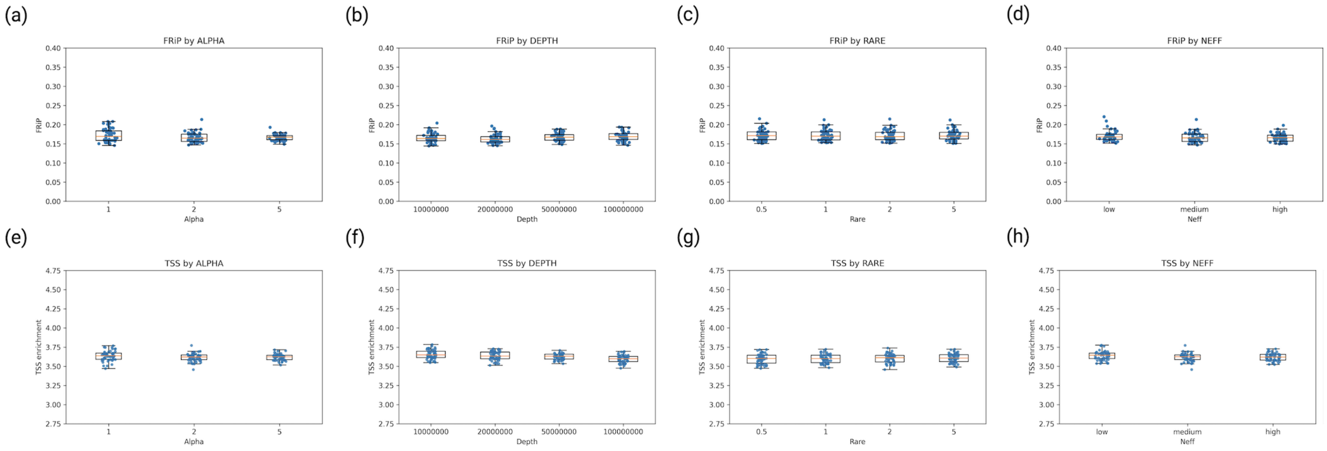


**Figure 2 Library quality control of the *in silico* pseudo-bulk panel.** Upper panels (a-d) show the fraction of reads in peaks (FRiP) and lower panels (e-h) show TSS enrichment scores, each stratified by (a,e) mixture evenness (Dirichlet $\alpha$), (b,f) sequencing depth, (c,g) rare-cell spike-in proportion, and (d,h) donor diversity ($N_{\text{eff}}$ bin).

We first verified that our method of generating the pseudo-bulk data varied these four characteristics appropriately (Figure 1). Evenness of cell proportions was parametrized using a Dirichlet distribution parameter ($\alpha = 1 < 2 < 5$), with lower $\alpha$ producing sparser, more imbalanced mixtures and higher $\alpha$ yielding more uniform ones. As shown in (Fig. 1a), the Shannon entropy of the target compositions increased monotonically with the Dirichlet concentration as expected. Second, we verified that the overall sequencing depth was accurate (Fig. 1b). We found that the ratio of observed to target reads centered near 1.0 with a mild negative bias and wider spread at 50-100M, attributable to alignment filtering and the ceiling introduced by downsampling. Third, we verified that simulated rare cell types were present at desired amounts (Fig. 1c), where the rare cell type fraction closely followed their target proportion (0.5-5%) with modest dispersion expected from per-cell read variability and filtering. Lastly, we verified that the desired number of samples was accurate based on a given range of either low (4-8), medium (12-16), or high (20-24) numbers of samples (Fig. 1d). These were summarized by the Simpson-effective sample number $N_{\text{eff}}$. Together, these checks confirmed that the synthetic data provided the desired range of characteristics in samples for downstream benchmarking.

### 2 Summary of WAD performance across wavelet decomposition levels and hepatic cell types.

Metrics were computed from 700 *in silico* pseudo-bulk realizations per condition. The "ALL" row represents the mean performance aggregated across six hepatic cell types (Cholangiocytes, Hepatocytes, Kupffer, LSEC, Mesenchymal, and NK-T).

| Wavelet level | Cell type | Mean MAE | Median MAE | IQR (MAE) | RMSE | Pearson $r$ |
|---|---|---|---|---|---|---|
| 1 | ALL | 0.0615 | 0.0507 | 0.0559 | 0.0783 | 0.85 |
| 1 | Cholangiocytes | 0.0648 | 0.0450 | 0.0669 | 0.0861 | 0.81 |
| 1 | Hepatocytes | 0.0779 | 0.0695 | 0.0452 | 0.0940 | 0.87 |
| 1 | Kupffer | 0.0509 | 0.0381 | 0.0469 | 0.0697 | 0.93 |
| 1 | LSEC | 0.0774 | 0.0666 | 0.0856 | 0.0977 | 0.61 |
| 1 | Mesenchymal | 0.0415 | 0.0341 | 0.0379 | 0.0530 | 0.89 |
| 1 | NK–T | 0.0566 | 0.0510 | 0.0530 | 0.0692 | 0.81 |
| 2 | ALL | 0.0638 | 0.0548 | 0.0633 | 0.0800 | 0.84 |
| 3 | ALL | 0.0636 | 0.0545 | 0.0631 | 0.0799 | 0.84 |
| 4 | ALL | 0.0635 | 0.0542 | 0.0627 | 0.0798 | 0.84 |
| 5 | ALL | 0.0633 | 0.0540 | 0.0627 | 0.0797 | 0.84 |

**Table 1. Performance summary of WAD across wavelet decomposition levels and hepatic cell types.** Each value represents the mean across 700 pseudo-bulk realizations.

### 3 TSS enrichment and chromatin QC

For each single-cell liver sample, transcription start site (TSS) enrichment scores were computed using *Signac* (v1.14.0). Fragment files were generated with *Sinto* (v0.9.0) from filtered, duplicate-removed BAM files. Promoter regions were defined as $\pm 2$ kb windows around annotated TSS coordinates from *EnsDb.Hsapiens.v86*, and TSS enrichment was calculated as the ratio between aggregate accessibility signal at promoter centers and mean signal within flanking regions.

| | | | | | |
|---|---|---|---|---|---|
| 151: 3.475 | 174: 3.678 | 331: 4.065 | 334: 4.01 | 342: 3.775 | 346: 3.632 |
| 348: 4.098 | 358: 3.921 | 360: 3.955 | 365: 4.116 | 366: 3.616 | 369: 3.285 |
| 374: 3.318 | 382: 3.351 | 435: 3.434 | 436: 3.583 | 440: 3.587 | 450: 3.707 |
| 457: 3.255 | 459: 3.645 | 465: 3.873 | 470: 3.637 | 485: 3.865 | 489: 3.541 |
| 618: 3.929 | 623: 3.677 | 627: 3.87 | 639: 3.939 | 662: 3.882 | 724: 4.192 |
| 733: 4.133 | 741: 4.073 | 751: 4.376 | 755: 4.104 | 767: 3.695 | 783: 3.455 |
| 786: 3.618 | 791: 3.643 | 794: 3.58 | | | |

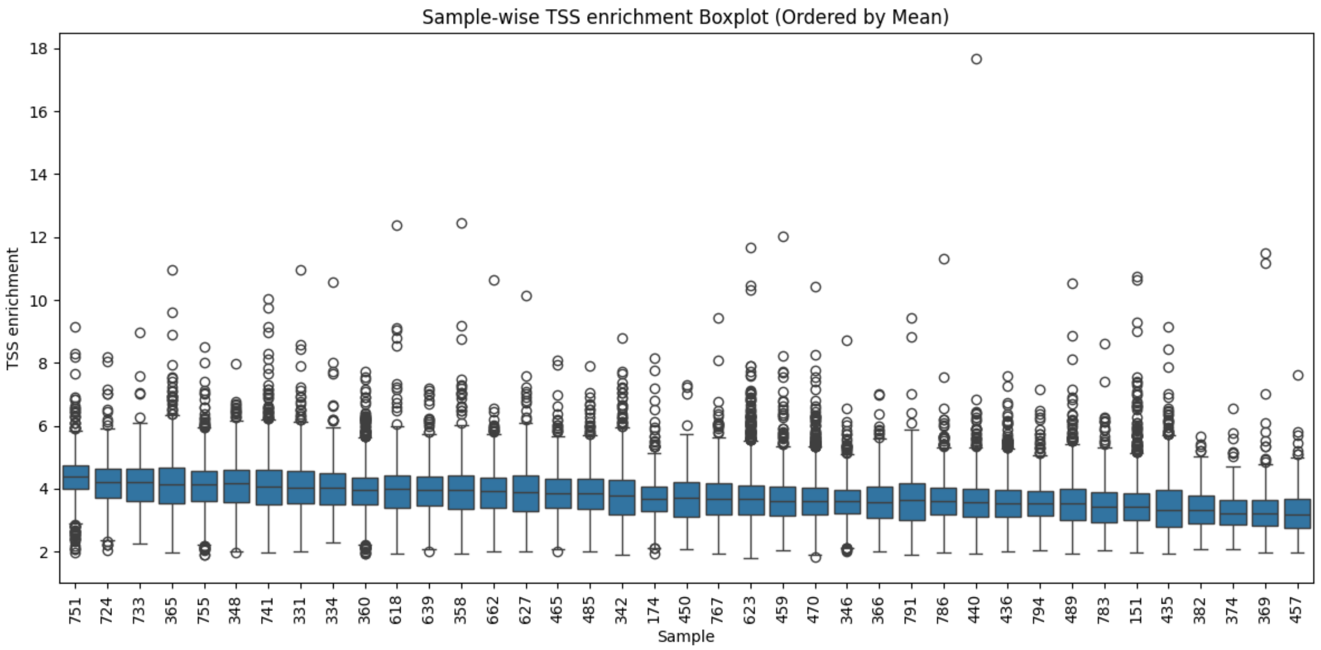


**Figure 1. Distribution of TSS enrichment scores across 39 liver samples.** Each box represents per-sample distribution, with circles indicating outlier barcodes. Most samples show consistent promoter-center accessibility (median $\approx 4$), indicating uniform library quality across samples.

### 4 BAM filtering and barcode selection

Cell-type–specific BAMs were generated by extracting reads with matching barcodes using samtools (v1.17). We restricted analysis to primary nuclear chromosomes (hg38) and applied stringent filtering to retain only high-quality, properly paired alignments. The exact command was:

```
samtools view \
  -h -b -f 3 -F 4 -F 8 -F 256 -F 2048 -F 1024 -q 30 \
  --tag-file CB:celltype_barcodes.txt \
  -L primaryNuclearChr.bed \
  -o {sample}_{celltype}.bam \
  {input_bam}
```

Resulting BAMs were sorted and indexed with:

```
samtools sort -o {sample}_{celltype}_sorted.bam {sample}_{celltype}.bam
samtools index {sample}_{celltype}_sorted.bam
```

### 5 Truth-prediction scatter plots

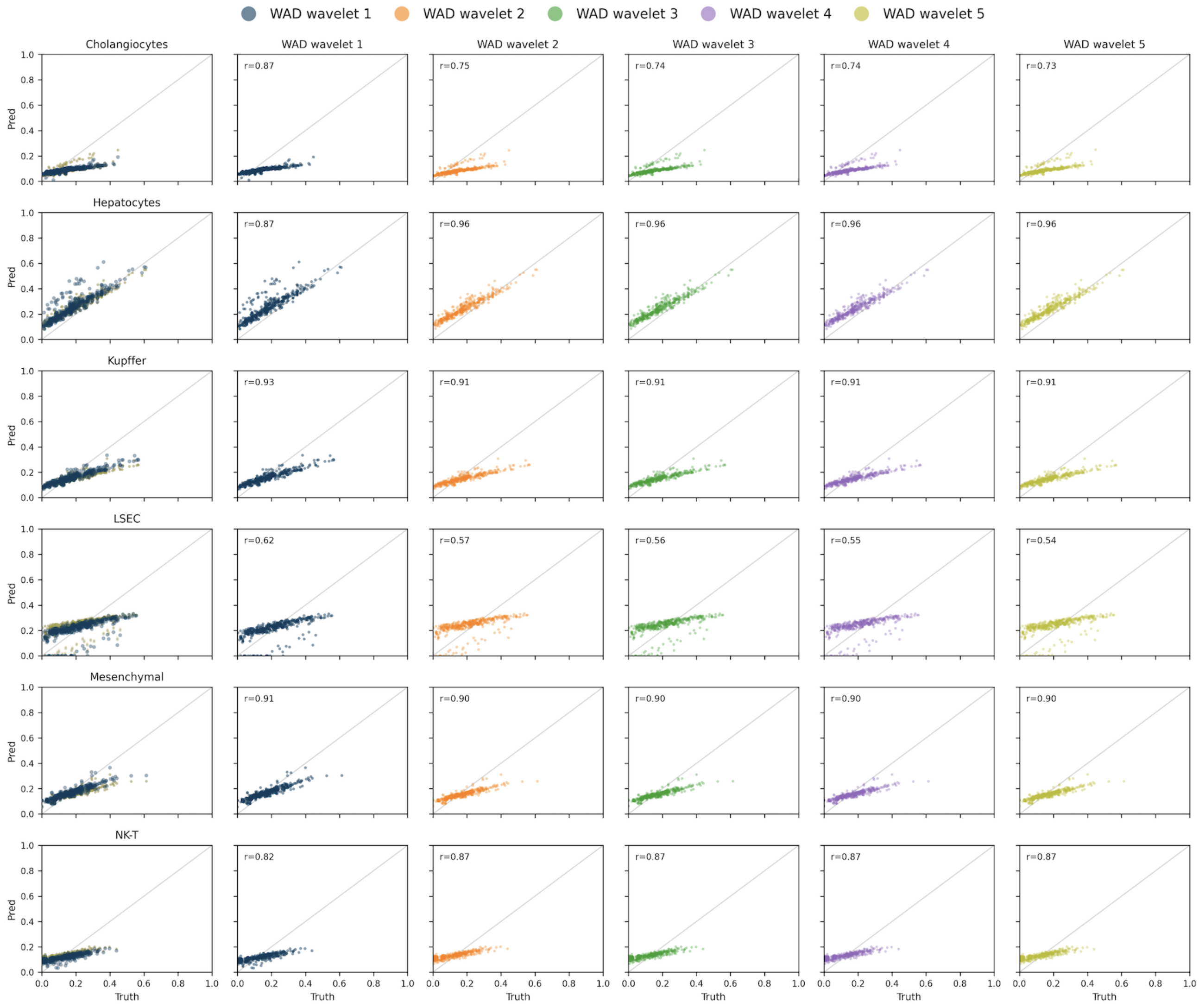


**Figure 1. Truth-prediction scatter plots across cell types for WAD wavelet levels.** Each point represents one *in silico* pseudo-bulk sample, with the x-axis showing ground-truth cell-type proportion and the y-axis showing the corresponding proportion. Rows correspond to cell types. The leftmost panel overlays all wavelet levels (1-5) for direct visual comparison, whereas the five panels to the right display individual wavelet levels separately. The grey diagonal line $(45°)$ denotes perfect agreement between predicted and true proportions. The Pearson correlation coefficient $(r)$ in each single-level panel quantifies concordance across all pseudo-bulk realizations for that cell type.
